# Quantum Chemical Profiling of Protein Mutations via Fragment-Based DFT

**DOI:** 10.64898/2025.12.17.694938

**Authors:** Alejandro Leyva, M. Khalid Khan Niazi

## Abstract

Missense mutations have been extensively studied in tumor-suppressing antigens (TP53) to understand oncogenesis within malignant epithelial cells. Using Whole Exome Sequencing (WXS), missense mutations can be profiled into protein sequences to identify the most common variants in tumor samples. Since nearly 80% of mutations arise randomly, it is necessary to isolate those that produce dysfunctional proteins within large cohorts. Using threading and generative algorithms such as AlphaFold and ColabFold, large cohorts of WXS information can be converted into computationally analyzable structures. By evaluating both high- and low-confidence regions in these structures, these antigens can be studied en masse using pipelines that generate analytical inputs for quantum chemistry analysis. we created a pipeline that processed whole-exome sequencing (WXS) data and selected 28 representative TP53 missense mutants from the TCGA-BRCA cohort for quantum-chemical feasibility analysis. These structures were systematically cleaned using tools such as OpenBabel and AmberTools, and each was prepared for Natural Population Analysis (NPA), Electrostatic Potential (ESP) calculations, and Highest and Lowest Occupied Molecular Orbital (HOMO/LUMO) evaluation within Q-Chem. Using this pipeline, population genomics can be integrated with chemoinformatics to analyze electron density concentrations and generate hypothesis-generating electronic descriptors associated with protein dysfunction. By modifying the generated inputs, additional analyses such as Fukui orbitals, chemical shifts, and Raman shifts can also be performed. This provides a computational means to probe electronic properties not readily accessible at scale using experimental techniques.

## 1 Introduction

The TP53 gene is a multi-domain transcription factor responsible for producing antigens that regulate apoptosis, cell cycles, autophagy, metabolism, and DNA repair [2,4]. Within the 393-amino-acid protein, DNA binding occurs through residues 92–292, producing upregulation or downregulation of specific cascades [1]. Nearly 90% of all mutations occur in this DNA-binding region, preventing regulation of the PIK3CA, Wnt, RAS, and ATM pathways [5]. Within this region, a beta-sandwich fold is induced through zinc coordination by Cys176, His179, Cys238, and Cys242 [2,3]. The loss of the metal ion relinquishes DNA-binding affinity, resulting in dysregulation of DNA-damage and repair pathways essential for mesenchymal plasticity within aggressive tumors [3].

The tetramerization domain in residues 323–356 is composed of a beta strand (326–333) followed by alpha-helical regions (335–355) that form a tetrameric structure connecting the domains of the p53 protein to form a dimer of dimers [6–9]. The hydrophobic core formed from isoleucine and leucine produces a packing interaction around the region, creating the coordinative ability for conformational changes sequential to DNA binding [6–9]. Threonine and glycine residues 329 and 334 form the bridging strand between the beta strand and the alpha-helical region. Arginine and lysine residues in positions 337 and 351 stabilize the salt bridge between asparagine and arginine that corrects the orientation of the alpha helix. The surface of the alpha helices is covered by hydrophobic residues including phenylalanine, leucine, isoleucine, and alanine. Orientation and parallelism of the helices are coordinated using isoleucine and leucine hydrogen bonds between amines. This produces a combination of van der Waals contacts between leucine and histidine while forming side chains between leucine and isoleucine, with glycine residues orienting the angular position of the helix at a 40-degree angle for symmetry. Certain mutations target the salt bridge or hydrophobic packing, preventing transcriptional activation and dimerization of p53 molecules [6–10].

The first 93 residues of p53 form transactivation domains 1 and 2 and the proline-rich region (62–93) [11–15]. TAD1 contains residues allowing for phosphorylation, while TAD2 contains short alpha-helical strands allowing transcriptional activation of PUMA, NOXA, and BAX [11–15]. The proline-rich region is polymorphic, containing PXXP motifs that enable protein binding. Finally, the C-terminal regulatory domain contains large clusters of lysines for ubiquitination, methylation, and acetylation.

Using whole-exome sequencing, extracted DNA from cell lines can be enzymatically fragmented and biotinylated, allowing identification of protein-coding regions [16–19]. Once extracted, these reads are processed in parallel to produce short reads that are compiled by protein region to assemble the amino acid sequence. Once the base-pair content is known, codons can be assembled to produce the full protein.

Typical experimental protocols synthesize mutated DNA by using restriction digestion to excise the protein region of interest [16–19]. Using synthetic DNA copies from phosphoramidite chemistry, a heteroduplex forms between mutant and wild-type sequences, and DNA polymerase synthesizes the mutant DNA, which is extracted using methylation-restriction digestion. Computationally, this process is far simpler, as the chemical behavior of proteins can be modeled using the sequence in AlphaFold with developed thresholding algorithms [20–24]. Using AlphaFold or ColabFold, multiple sequence alignments of protein inputs can be compared against evolutionary variants to predict protein geometry, which is then used to determine bond angles, turns, and secondary structures [20–24]. These generated protein structure files can be used for molecular dynamics simulations to analyze mechanistic behavior in isolated systems. However, since most mutations are random and functionally invariant, it is difficult to determine which p53 mutants are functionally significant.

Using Q-Chem, electron density distributions can be analyzed to identify electronic perturbations associated with protein mutations [25–29]. Q-Chem provides this analysis by solving the Schrödinger equation using density-functional theory and Gaussian basis functions, assuming electrons move within probability densities around nuclei. This can be used to understand transition states and activation states in proteins and the basis of chemical transitions within these systems. Using Natural Population Analysis, electron densities around nuclei can be quantified and compiled into natural orbitals, yielding stable means to measure charge and electron transfer. Electrostatic potentials and orbital occupations provide a way to track mechanistic differences within proteins for large-scale mutational studies. This provides a means to understand protein misfolding and altered reactivity within the tumor microenvironment. We hypothesize that AlphaFold can be used to generate large cohorts of PDB structures for quantum-chemical analysis for broad population-genetics studies [30].

## 2 Materials and Methods

Within the TCGA-BRCA cohort, 28 WXS samples were processed into FASTA files, with one control p53 PDB. The control TP53 sequence was derived from UniProt P04637 [30]. Each mutant was characterized by its specific substitution and documented. ColabFold was used to generate five biological replicates of each mutant protein structure [21–23]. Five ColabFold structural predictions were generated for each mutant; a single representative structure was selected for feasibility analysis to limit computational cost. OpenBabel was used to supply adequate hydrogens and ensure neutrality. Q-Chem 6.3.0 and Amber24 were used on the Ohio Supercluster’s Ascend cluster, running on NVIDIA100 GPUs [25–29]. AlphaFold 2.3.2 was accessed through ColabFold, in conjunction with Biopython, for structure inference [20–24]. All base-pair sequences for each mutant in the whole-exome dataset were stored in a CSV named after the file ID, converted into amino-acid labels, and written into FASTA files for AlphaFold generation.

RMSD values and chemical shifts were extracted using BioPDB, and all inputs for sequential Q-Chem analysis were generated using Biopython and BioPDB. All code is provided in the GitHub for accessibility. Each input was then run in Q-Chem using a sequential shell script, where Natural Population Analyses, Electrostatic Potentials, and orbital occupations (HOMO/LUMO) were computed [25–29]. Electrostatic potentials were analyzed using the following parameters:

**Table 1:** Q-Chem input parameters used for ESP charge calculation, NPA analysis, and HOMO/LUMO orbital extraction.

| **Parameter** | **ESP (CHELPG)** | **NPA** | **HOMO/LUMO** |
| --- | --- | --- | --- |
| JOBTYPE | SP | SP | SP |
| METHOD | B3LYP | B3LYP | B3LYP |
| BASIS | 6-31G* | 6-31G* | 6-31G* |
| SCF_CONVERGENCE | 7 | 7 | 7 |
| MAX_SCF_CYCLES | 200 | 200 | 200 |
| SCF_ALGORITHM | DIIS_GDM | DIIS_GDM | DIIS_GDM |
| SYM_IGNORE | 1 | 1 | 1 |
| NO_REORIENT | 1 | 1 | 1 |
| CHELPG | TRUE | – | – |
| NBO | – | 1 | – |
| PRINT_ORBITALS | – | – | TRUE |

**Figure 1:**
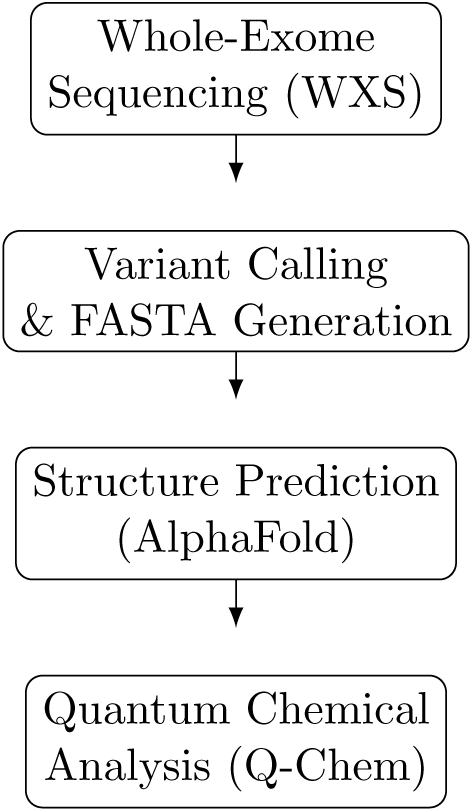
Compact multi-row flowchart summarizing the computational pipeline from WXS data to structural modeling (AlphaFold) and quantum chemical analysis (Q-Chem).

## 3 Results

The results for the sequence generation analysis and ColabFold generation show a low confidence within the proline-deep regions and a substantial portion of the DNA-binding domain, likely due to the absence of evolutionary variants [20–24]. This is reasonable, as most cancerous mutations are not natural or evolutionarily promoted [5]. In addition, there are more than 100 different mutations within the cohort that arise in patients with breast cancer, and these sequences originate from malignant cells. Since most mutations were synthetic, the MSA could not find an evolutionary match to the protein other than the original TP53. However, the protein generation was successful.

Analytically, most mutations occurred in the DNA-binding domain, which presents a statistically significant finding that these mutations arise despite the availability of residues [1,5]. Most mutations from this cohort were missense rather than nonsense mutations, which is statistically more likely based on prior literature on mutational patterns in TP53 [5]. Most protein mutations were non-truncated, and as expected, all nonsense mutations resulted in shorter proteins.

Biochemically, most of these mutations were evaluated to be non-conservative according to BLOSUM62 scoring and the recorded amino acid substitutions. Most mutations occur within residues 150–200, which correspond to the zinc-binding residues that functionalize TP53 DNA binding [1–3]. A secondary hotspot occurs between residues 250–300, corresponding to arginine-binding positions that orient the helical structures of the dimerization complex [1–3]. The highest-frequency mutation was the R157H mutation, which occurs in the DNA-binding domain and results in the downregulation of BM1 and increased expression of TWIST1, producing epigenetic deformation and unregulated gene expression [5]. Mutant variants such as R273H and R196, which have been found in myeloid acute leukemia and triple-negative breast cancer, respectively, correspond to worse prognosis [1–3,5]. The most common amino-acid substitution is from arginine to histidine, corresponding to the top mutations within the BRCA cohort, all of which destabilize the DNA-binding domain and contribute to proliferation and reduced survival [5]. The results for the wildtype alphafold generation found high confidence and stable generation of replicates, with evolutionary convergence and high confidence generations within the DNA binding domain. The predicted alignment error showed a high confidence across ranks. Wild-type generation of colabold stuctures was successful overall.

**Table 2:**
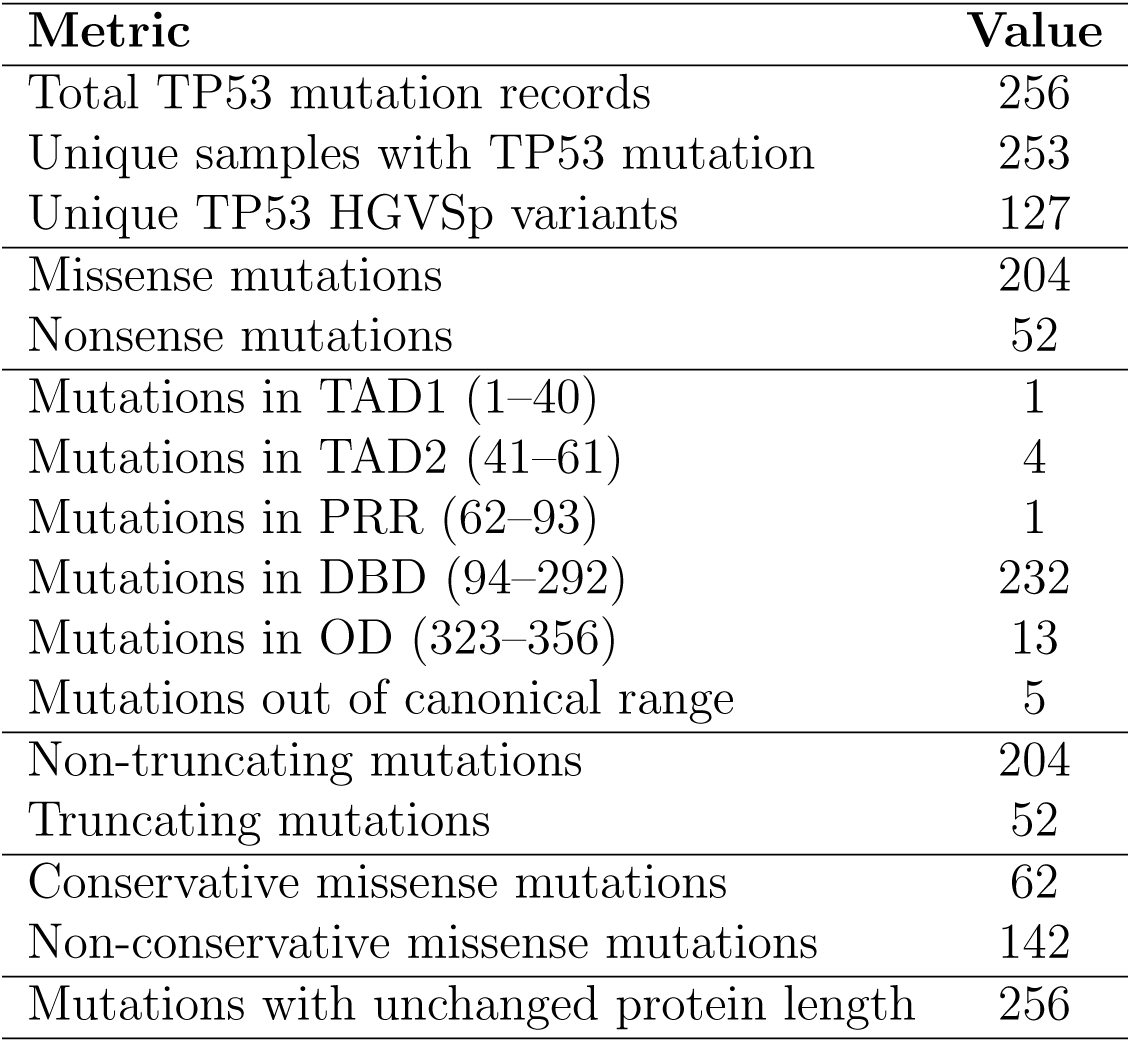
Summary of TP53 Mutation Statistics.

**Figure 2:**
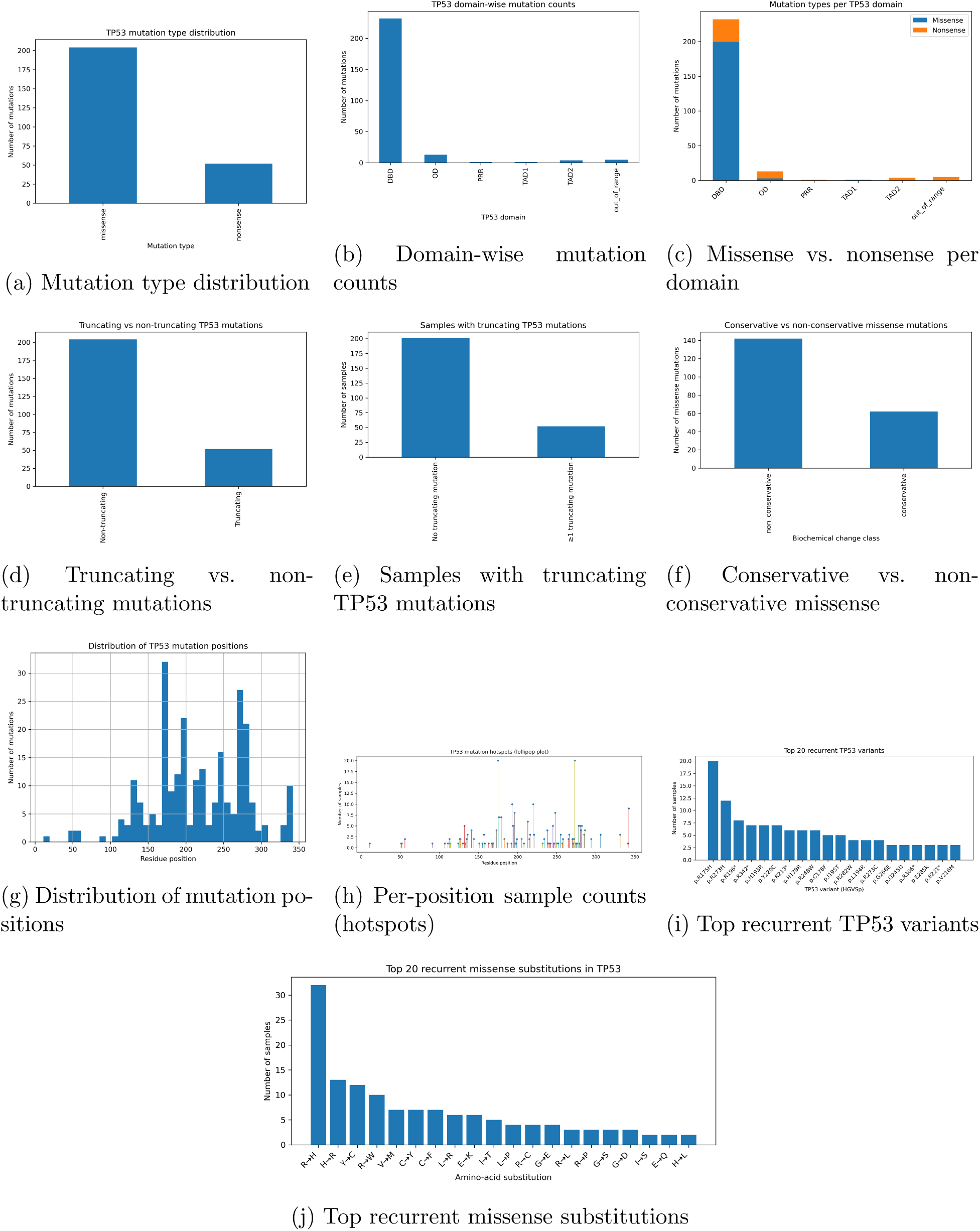
Comprehensive overview of the TP53 mutational landscape in the BRCA cohort. Panels summarize global mutation characteristics, truncation status and biochemical severity, positional distributions and hotspot structure, and the most recurrent TP53 variants and amino-acid substitutions.

**Figure 3:**
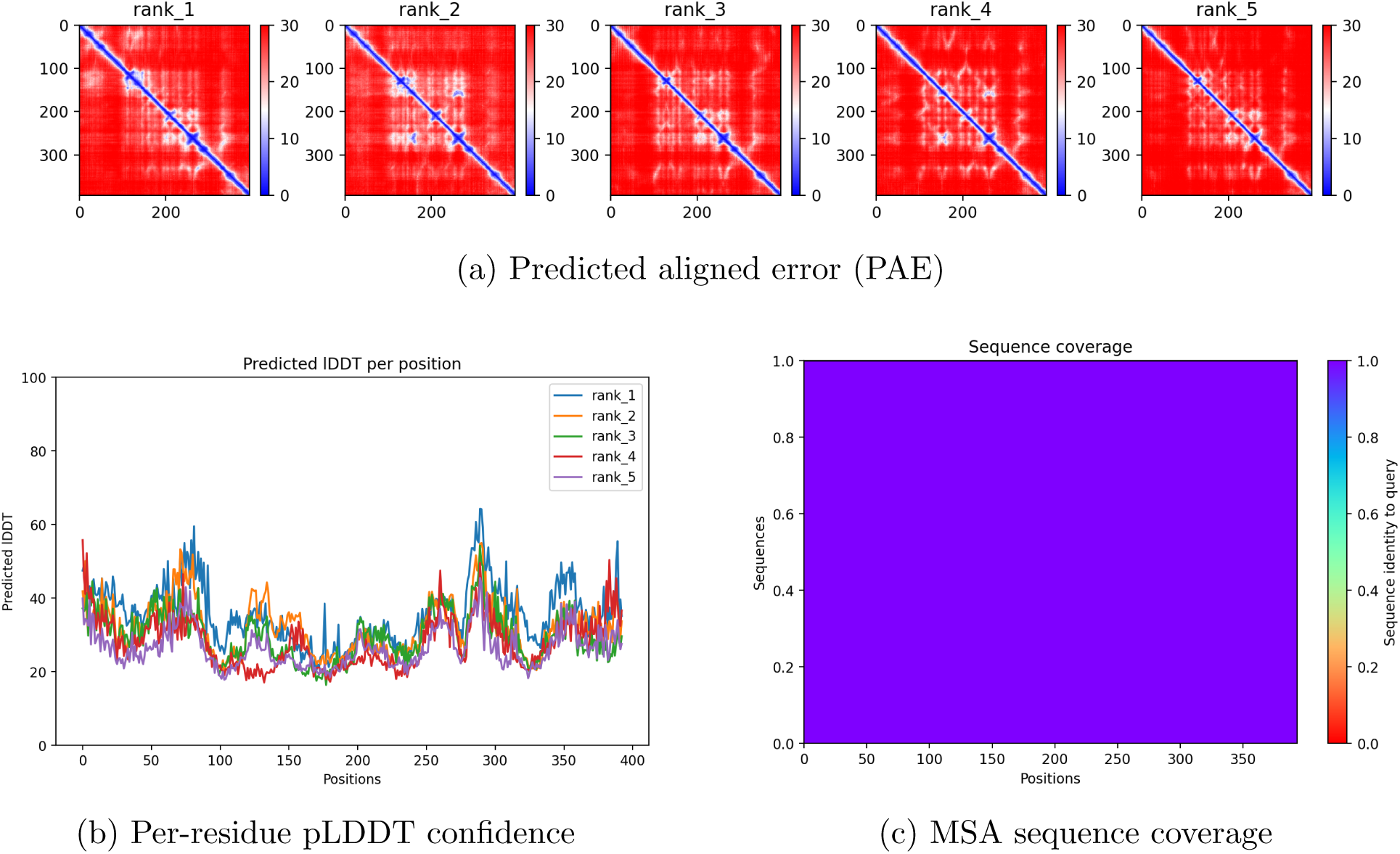
AlphaFold/ColabFold structural prediction outputs for TP53 C176 variants. The full-width panel shows the predicted aligned error (PAE). The bottom row shows per-residue pLDDT confidence scores and the MSA sequence coverage.

**Figure 4:**
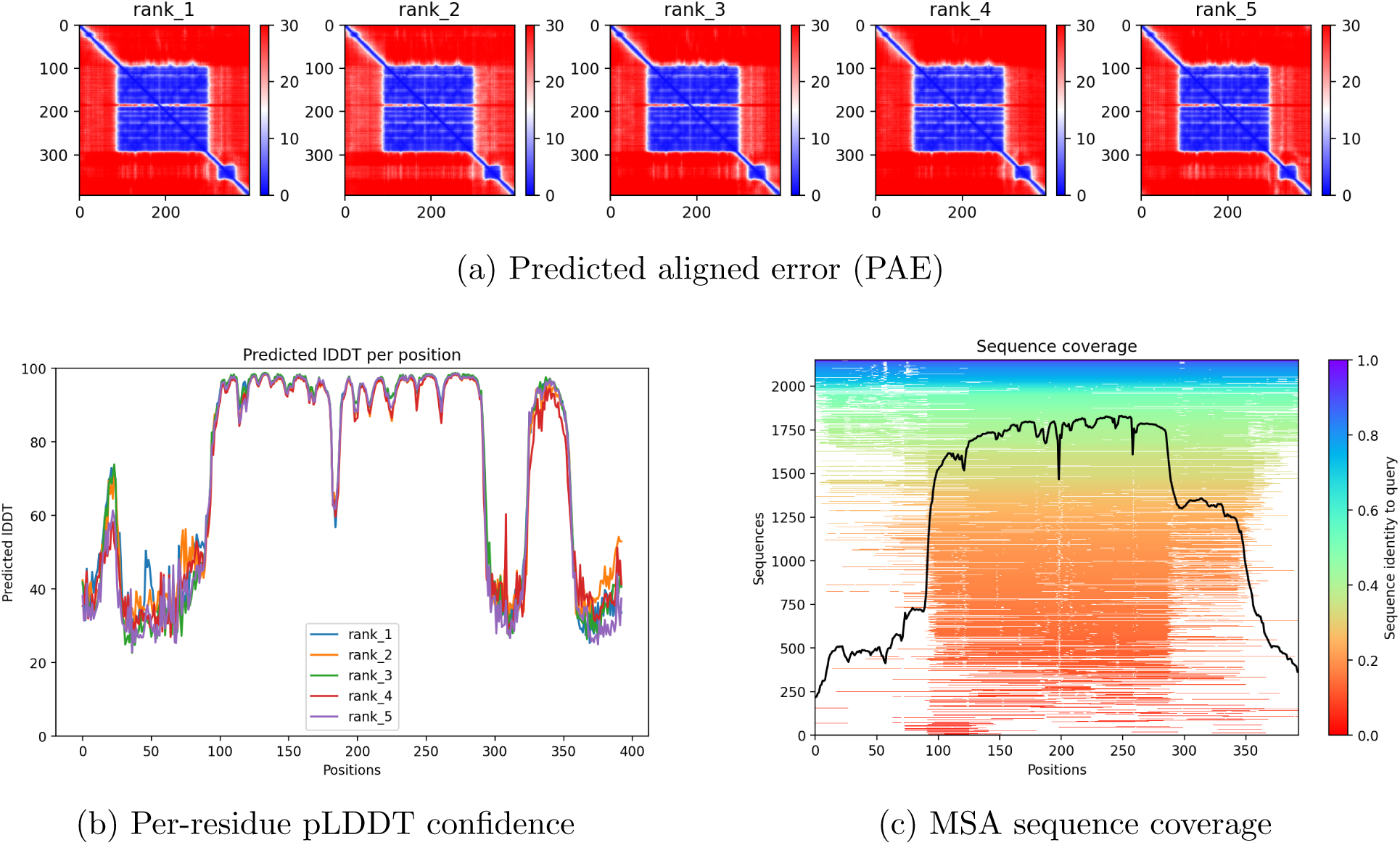
AlphaFold/ColabFold structural prediction outputs for TP53 Wild-Type The fullwidth panel shows the predicted aligned error (PAE). The bottom row shows per-residue pLDDT confidence scores and the MSA sequence coverage.

The display for the wildtype protein was extracted from the protein database bank, and the mutant structures were displayed on the left. Based on the generations, the beta sandwich folds were not conserved along with a substantial portion of the alpha-helical regions. low confidence regions are seen in the unstructured strands, and the low confidence wildtype strands are also shown within proline rich regions and transctivation domains. The structures were conserved adequately for Q-Chem analysis.

Across the 28 mutants, a lower electrostatic potential was observed across the cohort, characterized by substitution of polar amino acids in conserving domains in the DNA-binding region [1–3,25–29]. High variability was also demonstrated across all mutants, though most mutations resulting in neutral residue substitution had a similar variance. Mutants that resulted in substitutions of similar chemical characterizations showed similar variability. Within the mutant cohort were seven cysteine-mutated variants, all within the binding domain that sever the zinc linkage [1–3], one alanine mutant, three aspartate mutations, three glutamate mutations, three phenylalanine mutations, and three glycine mutants. Within this cohort, the glycine mutants typically substituted larger amino acids, while the glutamine mutants resulted in a complete shift from a negatively charged amino acid to a positive one. The phenylalanine mutants showed a shift toward polar thiols, representing a negative shift in charge, with most of these mutations concentrated within the dimerization complex [1–3,5]. The cysteine mutations showed a mixed set of potential shifts in both directions, while the aspartate mutations typically were substituted with neutral or hydrophobic amino acids. Across the atomic charge distributions within the fragments of the first five mutants, the cysteine mutants demonstrated an overwhelming similarity at the mutation point and showed correlation with large decreases in ESP [25–29]. The distribution across mutants shows a decrease in electric charge, which can be inferred from a subset of mutants—including glycine and cysteine variants—being replaced with polar residues. This infers that the polarity of the residue shifts the stability and binding capacity of the DNA-binding region [1–3,5].

**Figure 5:**
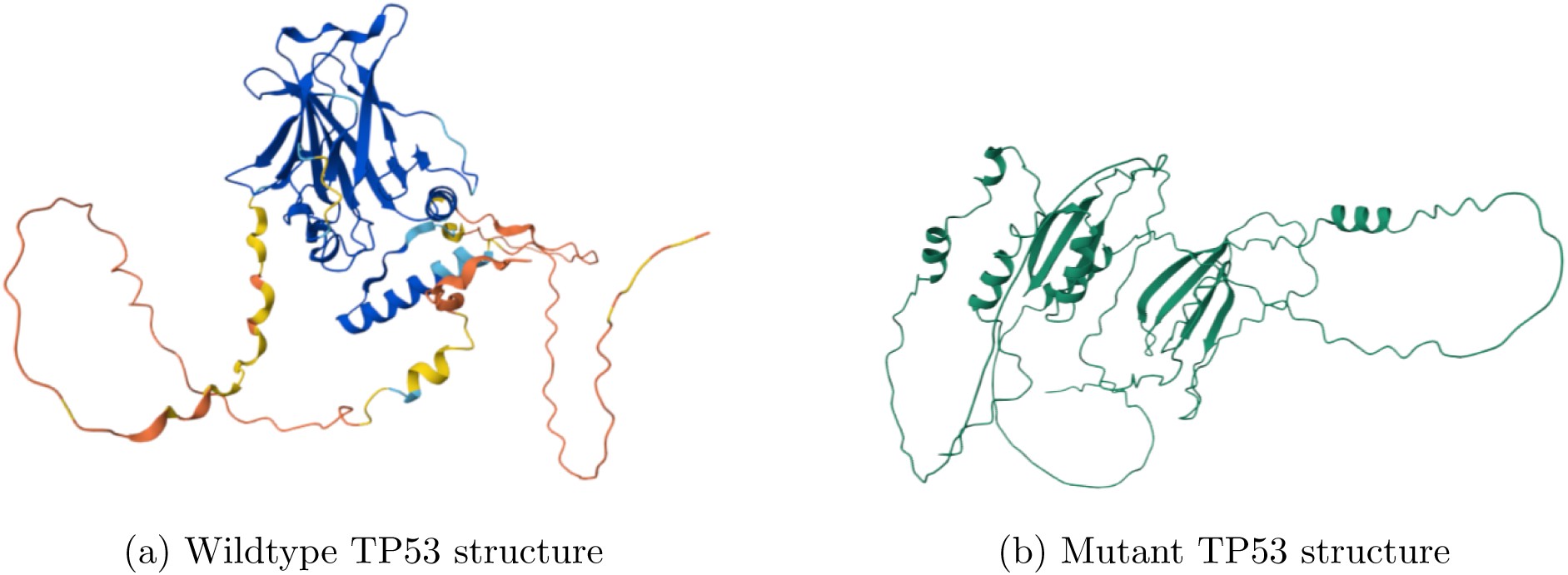
Comparison of the predicted structures for wildtype and mutant TP53. Structural differences are visualized using AlphaFold/ColabFold predicted coordinates.

Within the mutation fragments, the electronic distributions in Figure 8 were sparse. Across the first few mutants, the Natural Population Analysis demonstrated that each of the four cysteine mutations showed unique differences in charges across the atoms within the fragment, yet heavy convergence was noted on the third atom index, likely a component shared across all mutants [25–29]. The distribution of charge across the NPA output contained outliers, though a high frequency of observations between 0.4–0.5 when summed over atoms demonstrated high polarization within the fragment, with several mutants showing greater polarization approaching 0.7. The electron distributions in these mutants are overwhelmingly undersaturated, resulting in high polarization [25–29]. This is consistent with the mechanisms by which the orientation of the DNA-binding domain collapses under disrupted zinc coordination and altered charge density [1–3].

**Figure 6:**
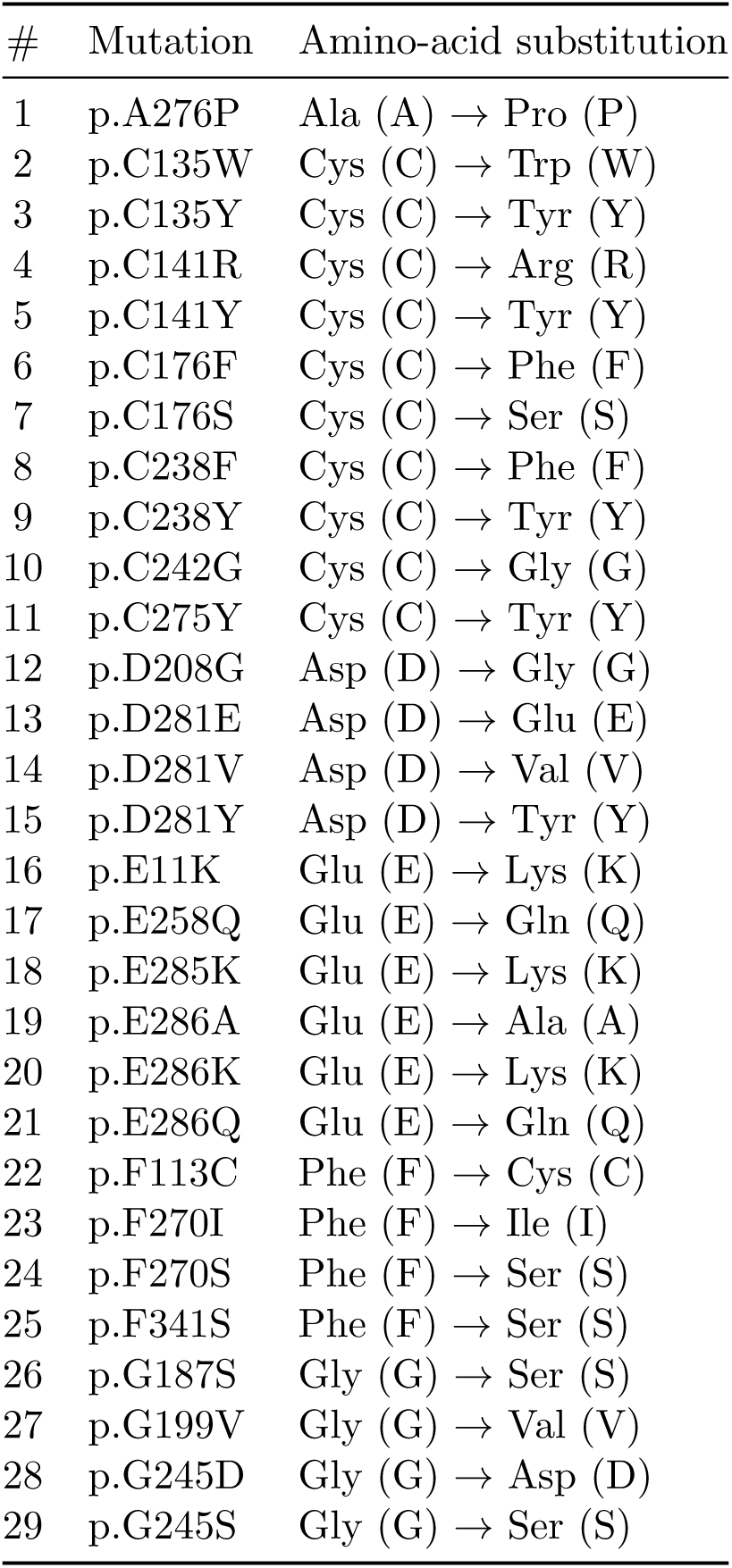
List of TP53 missense mutants analyzed in this study, showing residue position and amino-acid substitution relative to the wild-type sequence.

**Figure 7:**
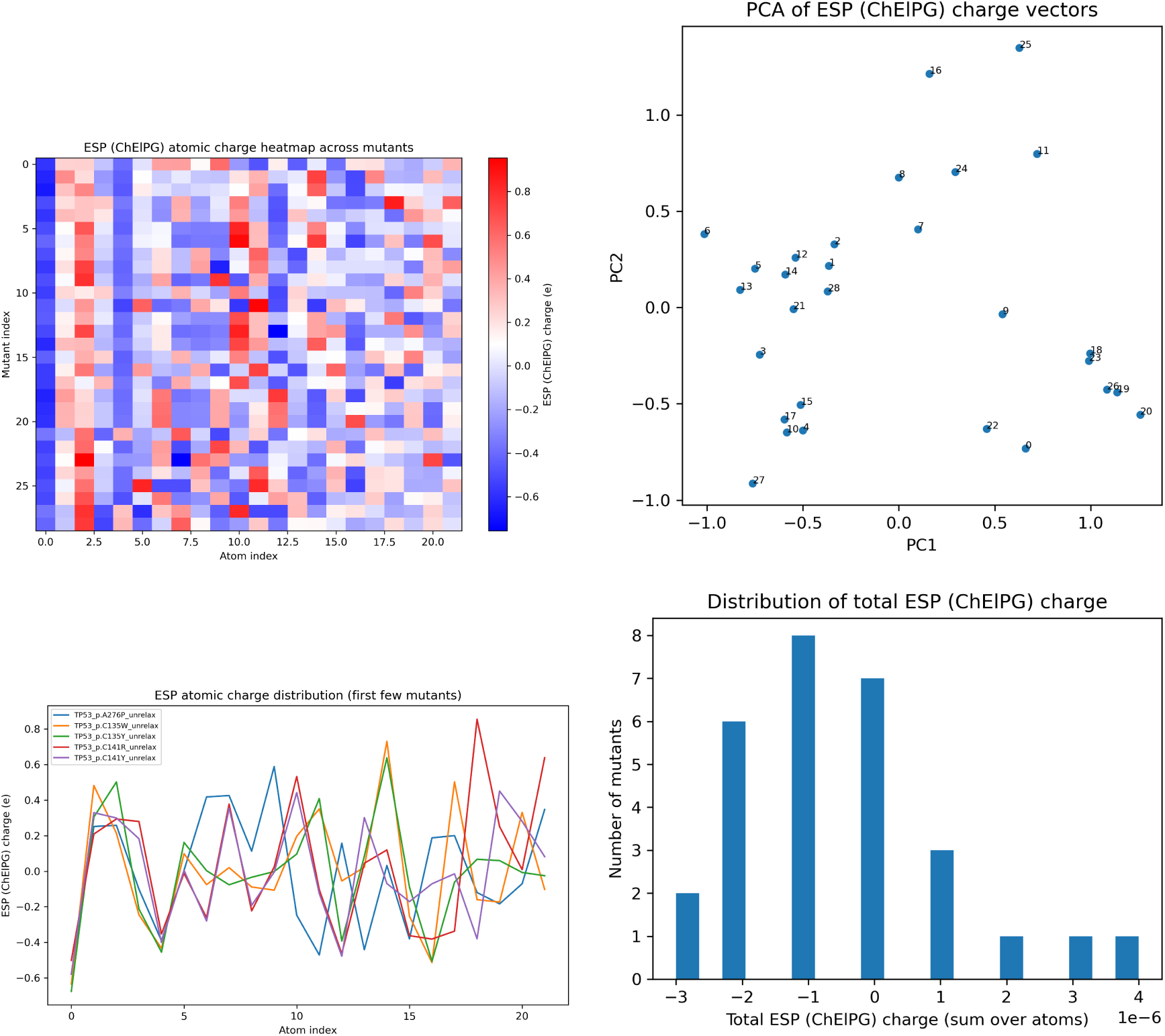
Electrostatic potential (ESP) analyses including heatmap, PCA, first-mutant line plots, and histogram of summed ESP distributions.

**Figure 8:**
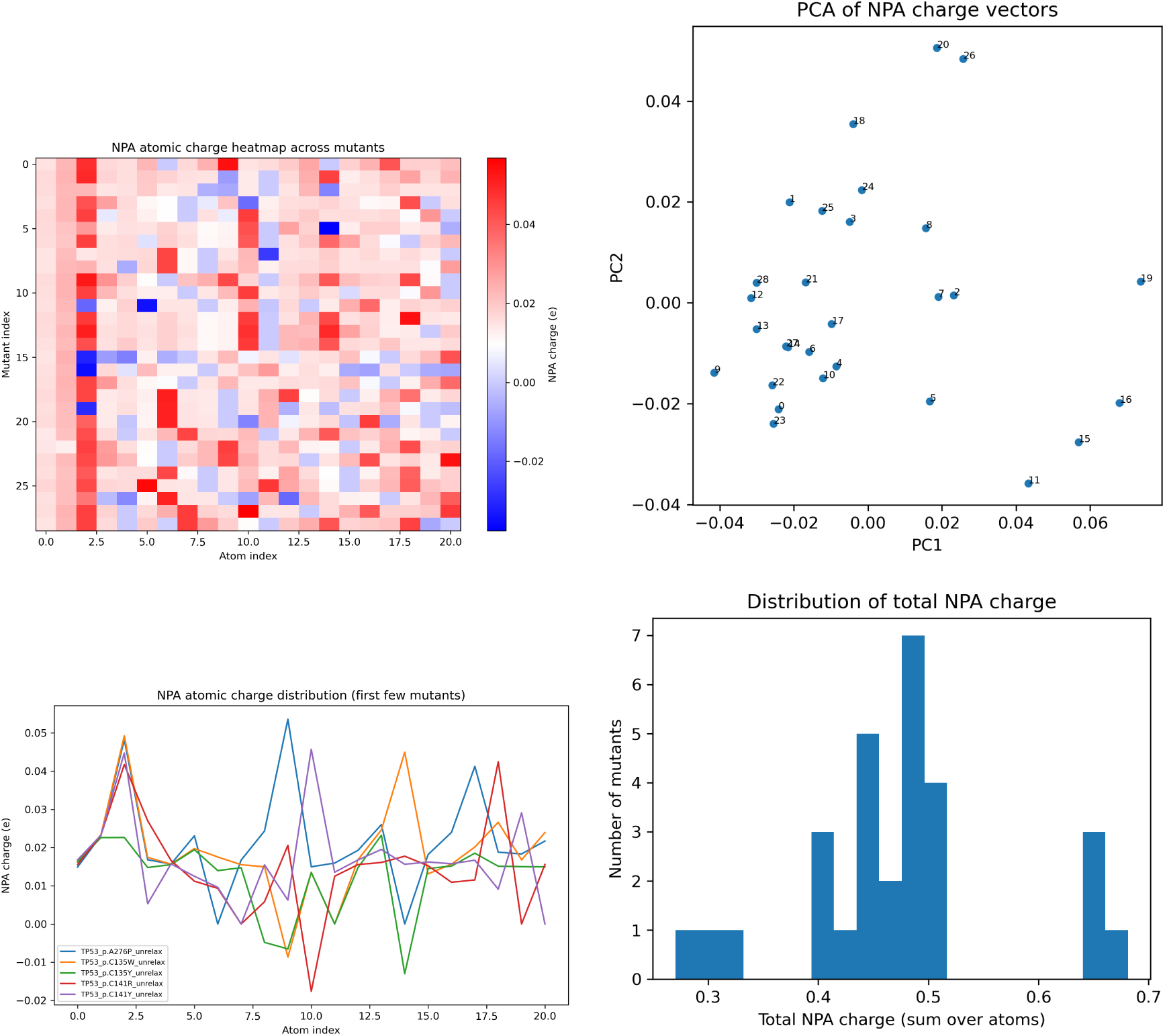
Natural Population Analysis (NPA) charge analyses including heatmap, PCA, first-mutant line plots, and histogram of summed NPA charges.

Across the HOMO and LUMO distribution in figure 9, a high frequency of mutants displayed an overwhelmingly electronicaly stable distribution, which presents a limitation of the analysis. While ESP and NPA results conclude that the electrostatic potential is reduced and that the electron distribution across fragments are sparse, HOMO/LUMO analysis indicated relatively stable frontier orbital energies across mutants, underscoring the limited discriminatory power of HOMO–LUMO metrics in this context. Many Oribtal energies were much lower than -1.5, suggesting a higher electronegativity within these atomic regions, though it is apparent that across samples, the electronic stability across atoms is consistent within all mutants.

**Figure 9:**
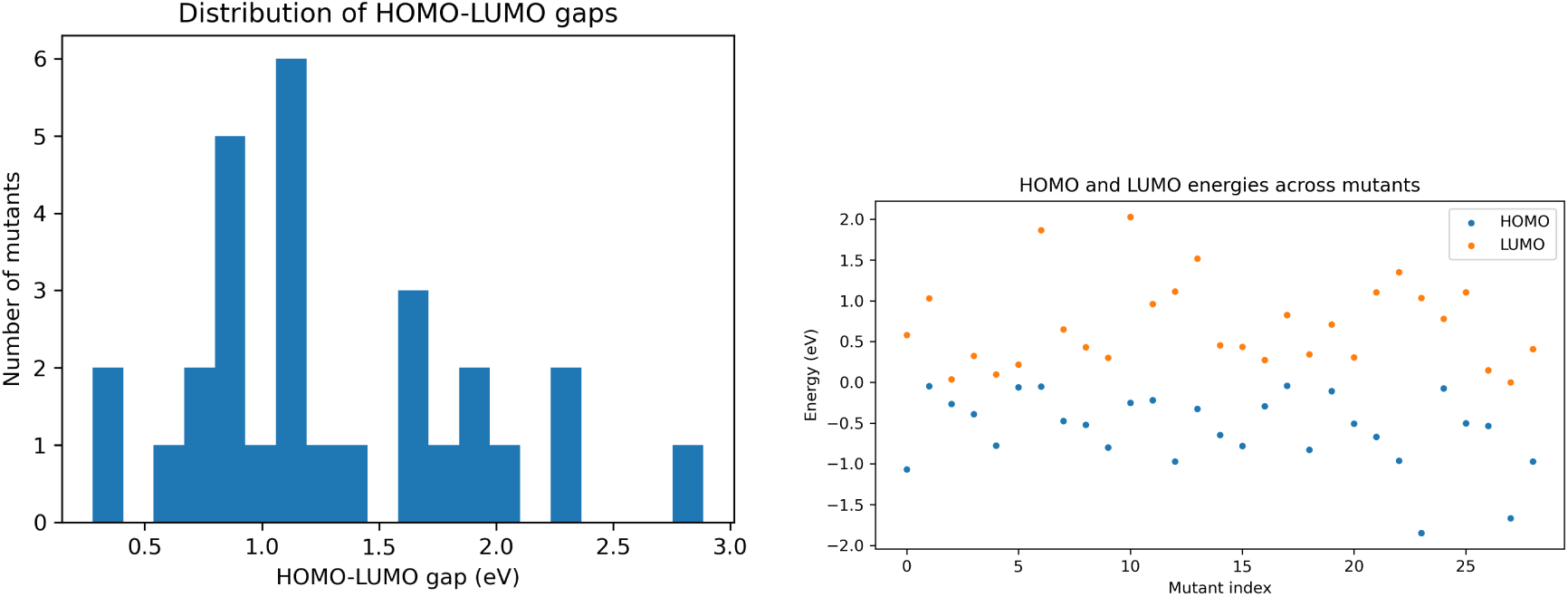
Quantum mechanical electronic structure metrics including HOMO–LUMO energy gap histogram and HOMO–LUMO scatter distribution across mutants.

In order to compare the unstable mutants to the wild type, the differences between the quantum electronic properties of each mutant and the wild type were graphed for NPA, ESP, and HOMO/LUMO analysis in Figure 10 [25–29]. Across each atom in the mutant structures, there is a consistently higher electronic sparsity as compared to the wild type, with the exceptions of indices 10 and 20, likely corresponding to hydrogenous regions. The ESP values for the mutants were lower than the wild type across most variants. This suggests that electronic sparsity and reduced electrostatic potential may contribute to destabilization of DNA-binding or tetramerization regions by decreasing the electronegativity in these regions [1–3]. The orbital energies were much higher in the mutants than in the wild type, indicating lower electronegativity across the mutants [25–29]. While these conclusions are limited by low-confidence ColabFold generations, they remain consistent with prior literature on mechanisms of TP53 affinity and structural destabilization [1–3].

**Figure 10:**
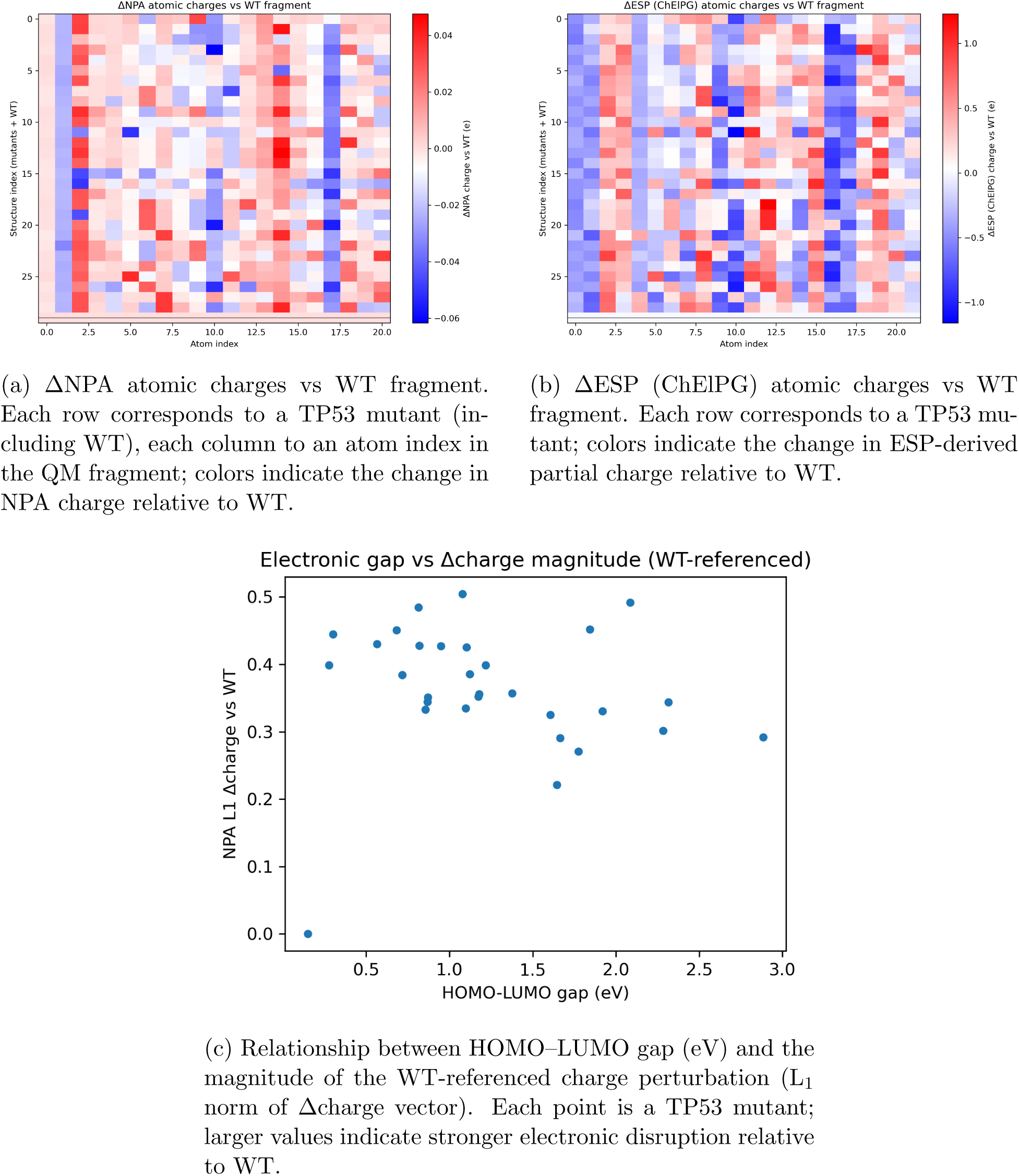
Quantum-chemical perturbation of TP53 mutants relative to the wild-type (WT) fragment. (a) and (b) show WT-referenced changes in NPA and ESP atomic charges, respectively, highlighting residues and environments with strong electronic redistribution. (c) links frontier-orbital perturbation (HOMO–LUMO gap) to the total magnitude of charge reorganization, summarizing how severely each mutant distorts the local electronic structure relative to WT.

The study presents a successful synthesis of mutant pdb forms, with electronic patterns characterized by lower electronegativity and sparse charge distribution. Likewise, there is lower electrostatic potential in these mutant types, reflecting low sample variability.

## 4 Discussion

Population level mutant analysis is possible using mass analysis of WXS sequencing. Transfer of these informatics into Fasta and sequential ColabFold was successful, though confidence is unremarkable. Electrostatic potential was consitently lower in the mutant cohort due to low sample size, and the electronic distribution was consistently more sparse, likely due to heavy residues and mutations with aromatic groups. Nevertheless, These mutations consistently exhibited lower electronegativity, which is compatible with previously reported destabilization mechanisms of DNA binding domains and tetramerization regions. Orbital analyses demonsrated higher electronic stability within the mutants, and more analysis can be performed to produce Raman spectra, chemical shifts, and nuclear magnetic resonance across these fragments. These analyses can be used to perform cohort level quantum characterization of gene mutations crucial in oncogenesis, more specifically, oncogenic processes. Mass analysis of cancer cohort data can provide general mechanistic observations and characteristics as to how mutations result in dysfunction of proteins. We generated structures for 28 mutants within the BRCA cohort using a developed pipleine and performed large scale quantum chemical analysis to understand the general trends for mutative behavior in TP53. For future studies, more features can be developed specifically for high confidence mutation prediction, in conjunctionn with physiochemical analysis.

## 5 Limitations

This study is limited by the low confidence generation of ColabFold, and does not use all 5 biological replicates. While the Q-Chem analyses are complete, the limited sample size constrains statistical generalization, positioning this study as a feasibility analysis. The pipeline is also limited by the use of ColabFold generated wildtype. MSA was not able to generate high confidence evolutionarily similar proteins for prediction of high confidence regions, such that in depth analysis of mutations in other domains is limited. Limited sample size of the BRCA cohort means analysis of mutations are validative in respect of replicating previous observations in prior studies.

## 6 Ethics Statement

All datasets are public and anonymized.

## 7 Author Contributions

All work was performed by the first author with supervision by the second author.

## 9 Data Availability

The datasets and diagnostic slides are publically available and anonymized. All code is available on GitHub.

## 10 Declaration of generative AI

During the preparation of this work the authors used ChatGPT in order to correct grammar and punctuation. After using this tool/service, the authors reviewed and edited the content as needed and take full responsibility for the content of the published article.

